# A scalable screening platform for phenotypic subtyping of ALS patient-derived fibroblasts

**DOI:** 10.1101/2022.09.27.509770

**Authors:** Karl Kumbier, Maike Roth, Zizheng Li, Julia Lazzari-Dean, Christopher Waters, Ping Huang, Vlad Korobeynikov, New York Genome Center ALS Consortium, Hemali Phatnani, Neil Shneider, Matthew P. Jacobson, Lani Wu, Steven Altschuler

## Abstract

A major challenge for understanding and treating Amyotrophic Lateral Sclerosis (ALS) is that most patients have no known genetic cause. Even within defined genetic subtypes, patients display considerable clinical heterogeneity. It is unclear how to identify subsets of ALS patients that share common molecular dysregulation or could respond similarly to treatment. Here, we developed a scalable microscopy and machine learning platform to phenotypically subtype readily available, primary patient-derived fibroblasts. Application of our platform identified robust signatures for the genetic subtype FUS-ALS, allowing cell lines to be scored along a spectrum from FUS-ALS to non-ALS. Our FUS-ALS phenotypic score negatively correlates with age of diagnosis and provides information that is distinct from transcript profiling. Interestingly, the FUS-ALS phenotypic score can be used to identify sporadic patient fibroblasts that have consistent pathway dysregulation with FUS-ALS. Further, we showcase how the score can be used to evaluate the effects of ASO treatment on patient fibroblasts. Our platform provides an approach to move from genetic to phenotypic subtyping and a first step towards rational selection of patient subpopulations for targeted therapies.

## INTRODUCTION

Amyotrophic Lateral Sclerosis (ALS) is an idiopathic, rare, and fatal neurodegenerative disease that displays considerable clinical and genetic variability^1^. Over 25 genes have been linked to ALS, each associated with a range of disease-causing mutations^2^, and nearly 90% of cases occur with no identified genetic cause^3^. Currently there are few treatments and no cure for ALS^4^.

From the perspective of developing therapeutics, it is unclear how to stratify ALS patients into subgroups that would respond similarly to treatment. Genetics provides a core framework for understanding the molecular basis of ALS^5–13^. Yet, even among individuals with the same disease-causing mutation, ALS shows variability with respect to age of onset, location of onset, cognitive changes, and disease progression^14–16^. mRNA profiling in cells and tissue has also been used to study ALS^17–20^ and to identify subgroups within a cohort including patients without known ALS genetic mutations^21^. However, mRNA alone may miss disease-relevant signals as ALS is also known to affect spatial organization at cell level, such as changes to protein localization and aggregation^22^. High-content microscopy can capture these spatial signals and has been applied in the context of Parkinson’s disease to classify disease vs. health ^23^ over a library of patient-derived fibroblasts. The use of fibroblasts is compelling, as fibroblasts are readily obtainable from living donors, retain the age of the patient, and can be scaled for high-throughput experimental assays. We hypothesized that primary patient-derived fibroblasts may provide disease-relevant information in ALS^24^.

We developed a platform to phenotypically profile primary ALS fibroblast cell lines, combining high-content imaging and machine learning to define a phenotypic score that measures a “distance” of each cell line to non-ALS control (“WT”), thereby quantifying a disease spectrum. Our phenotypic scores, based on immunofluorescent imaging markers of Early Endosome Antigen 1 (EEA1) and FUsed in Sarcoma (FUS), revealed strong separation between FUS-ALS and WT. The FUS-ALS phenotypic score negatively correlates with age of diagnosis and provided information that is distinct from transcript profiling. We show that this score can be used to identify sporadic patient fibroblasts that have consistent pathway dysregulation with FUS-ALS. Finally, we showcase how the score can be used to evaluate ASO treatment effects on patient fibroblasts.

## RESULTS

Establishing a scalable, image- and machine learning-based platform to identify phenotypes that separate ALS subtypes from WT patient-derived fibroblasts required the selection and optimization of three interrelated parameters: cell lines, molecular readouts, and image-derived features. To maximize biological interpretability, we sought separation based on a small number of cellular features and imaging markers related to known ALS pathology. With these requirements, we developed a robust screening protocol and machine learning pipeline to iteratively search for separation across a broad spectrum of ALS patient subtypes and library of imaging markers (Fig. 1a).

**Figure 1:**
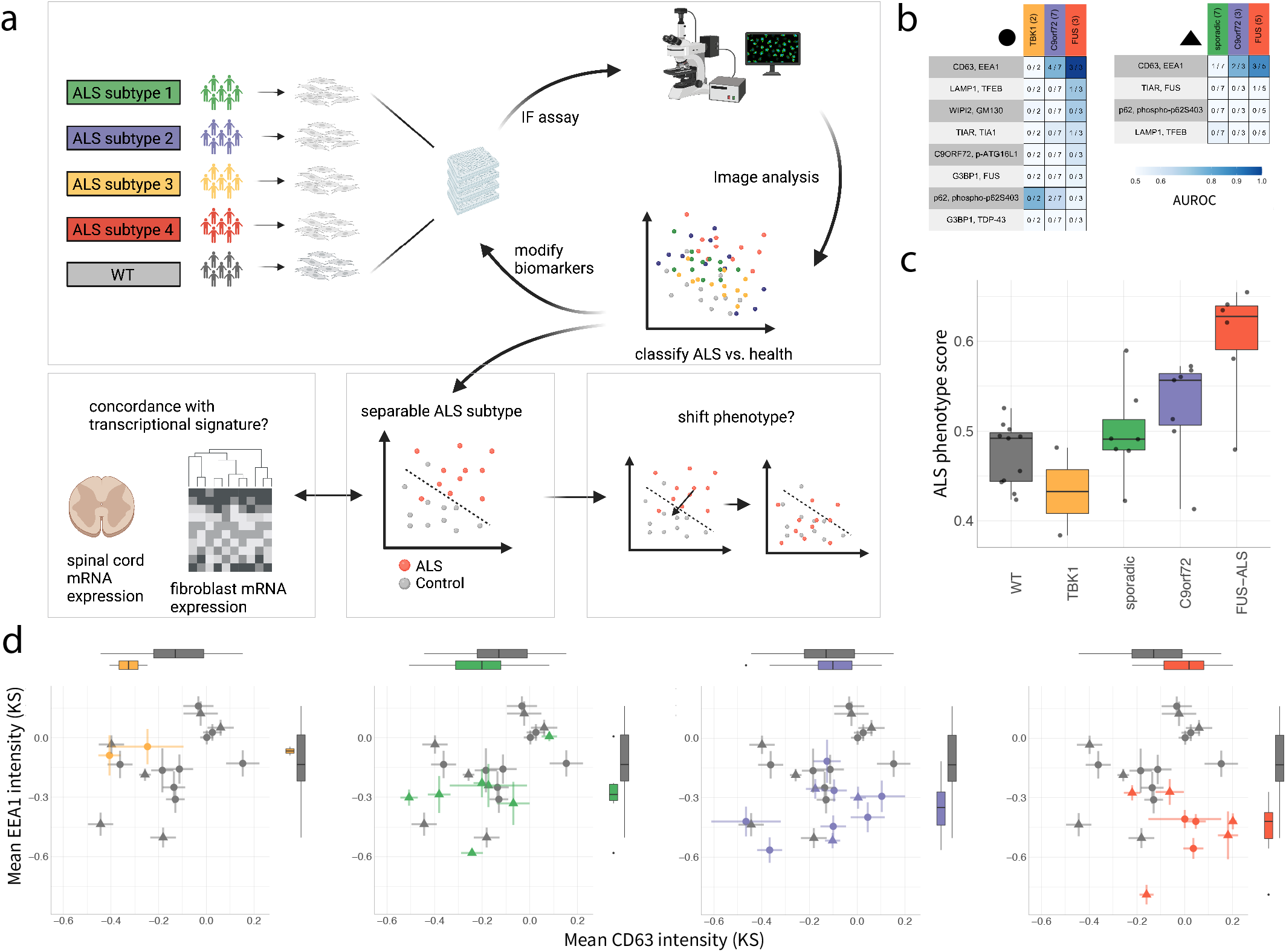
Overview of ALS phenotypic profiling platform and iterative search for biomarkers and ALS subtypes. **a,** Cartoon indicating iterative search strategy (top) and use cases in study (below). **b,** Classification accuracy of various ALS subtypes based on different combinations of biomarkers. Entries: fraction of cell lines separable from WT; color: area under the ROC curve (AUROC) evaluated across well replicates. **c,** Average ALS phenotypic score per cell line. Dots: cell lines. **d,** Two-dimensional projection of ALS subtype (colored) and WT (gray) profiles. Axes: best performing biomarkers/features for FUS-ALS subtype. Dots: averaged Kolmogorov-Smirnov (KS) profiles over well replicates and screen batches; KS relative to a WT cell line (Methods); shapes: screen batch as in (b); lines: 1 standard deviation of KS profiles. (b-d) Colors coded to ALS subtypes in b. (c-d) Box plots: median (center line), interquartile range (box) and data range (whiskers).

We first collected a cell line library of patient-derived fibroblasts that encompassed a variety of ALS backgrounds, including patients with mutations in TBK1, C9orf72, and FUS, as well as “sporadic” patients with no known disease-causing mutations and healthy (WT) controls (supplementary Table 1). Cell lines within each disease background were diverse with respect to specific mutations, age distribution, and sex (supplementary Table 1, Supp Mat). To measure cellular phenotypes, we surveyed a library of imaging markers using high-content microscopy. In total, we considered 13 imaging markers, assayed in multiplexed sets of 4 markers (supplementary Table 2). Every tested marker set contained two constant markers for automated identification of cellular (Mitotracker) and nuclear (Hoechst) regions. The other two variable markers measured a range of biological processes proposed to be related to ALS disease pathology, including autophagy, endo-lysosomal trafficking, and RNA processing. Applying these imaging markers to our collection of cell lines allowed us to assess patient-patient variability across a wide range of biological processes both within and between genetic backgrounds.

We optimized the screening platform to minimize experimental variability. In brief (Methods): all cell lines were grown for three passages to ensure full recovery from cryopreservation. Seeding density was optimized in 384-well imaging plates for automated image segmentation. For each batch and marker set, all compared cell lines were seeded in replicate plates, with a full set of all cell lines present on each plate. Finally, robotics was used to minimize technical variation in cell seeding and immunofluorescence labeling. These steps helped limit technical variability and highlight biological, disease-relevant differences between ALS subtypes vs. WT cell lines.

To profile each cell line, wells were initially imaged at 20X magnification and individual cells identified. Then, for each cell, ~200 image-derived features were quantified from the two variable markers (e.g., protein intensity, localization, and aggregation) (Methods). Using these features as predictors, a series of iterative random forest (iRF) classifiers^25^ were trained to discriminate between ALS and WT cells (Methods). This modeling choice has the benefits of: (i) imposing sparsity constraints that encourage models to make predictions from a small set of features; (ii) providing clear measures of each feature’s contribution to model predictions, measured as the mean decrease in Gini impurity (MDI); and (iii) allowing for hierarchical groupings of ALS vs. WT, where each group can have within-group heterogeneity (e.g., across different ALS subtypes, cell lines within each subtype, individual cells within one cell line)^25,26^. We reasoned that these models could capture biological heterogeneity and identify interpretable signatures that differentiate ALS from WT cells.

Each cell was scored according to iRF model predictions, and scores were averaged within each cell line (in comparisons of cell lines) or well replicate (in comparisons of well replicates). Prediction scores for a given cell line were generated from models fit with that cell line held-out from training. Thus, prediction scores directly assessed the generalizability of learned phenotypes to “unobserved” cell lines. To evaluate the degree to which different ALS genetic backgrounds correspond to distinct ALS subtypes, the distribution of predictions scores between each genetic background and WT were compared (Methods).

We first sought to identify imaging markers capable of separating any ALS background from WT. Six of the nine imaging marker sets provided no meaningful separation. However, three of the nine tested imaging marker combinations separated at least one FUS-ALS cell line from WT (3 FUS lines, 9 WT lines; Table 1) and two of nine separated at least one C9Orf72 cell line from WT (Table 1; Fig. 1b). Thus, we expanded our cell line collection for FUS-ALS (+3) and WT (+2), and we added 7 sporadic-ALS (supplementary Table 1). Consistently, the strongest separation was observed for the FUS-ALS subtype using imaging markers associated with intracellular vesicles, CD63 and Early Endosome Antigen 1 (EEA1) (Fig. 1b). This separation was primarily driven by the intensity ratios of EEA1/CD63 and the intensity level of EEA1 within the cell (Gini impurity feature importance; Methods). These features also showed notable, though weaker separation for C9orf72 (recall C9orf72 = 4/7 vs. recall FUS-ALS 5/6; Fig. 1c). Based on the separation scores (Fig. 1c-d), we chose to follow up with FUS-ALS.

We tested whether the observed FUS-ALS vs. WT separation would generalize to new cell lines with FUS mutations. Our initial screens included a small number of FUS-ALS samples—FUS-ALS comprises a small percent (~2-5%) of familial ALS ^27^. We acquired and included an additional 6 FUS-ALS cell lines (supplementary Table 1). With our focus on FUS-ALS, we also revisited the possibility of using an imaging marker for the FUS protein itself. The FUS imaging marker showed weak separation in our initial screens (Fig. 1b). However, prior work demonstrated that FUS shows cellular phenotypic differences between WT and FUS-ALS in isogenic iPSC lines^28,29^, in motor neurons differentiated from human patient-derived fibroblasts^30^, postmortem tissue^31^ and mouse spinal cord^32^. Given our expanded repertoire of FUS-ALS cell lines, we hypothesized that the FUS imaging marker would improve separation performance relative to our optimized EEA1/CD63 imaging marker set. We compared all pairwise combinations of the FUS, EEA1, and CD63 imaging markers across 12 FUS-ALS and 11 WT cell lines. Pairs including the FUS imaging marker improved separation relative to EEA1/CD63 (Extended Data Fig. 1). While the CD63/FUS marker set slightly outperformed the EEA1/FUS marker set relative to cell line classification, separation from the EEA1/FUS marker set was more robust across well replicates and single cell classification. For subsequent studies, we chose to use EEA1/FUS as our primary choice for classification together with Hoechst/Mitotracker for nuclear/cell identification.

FUS-ALS tends to be early-onset, and the distribution of FUS-ALS cell lines in our previous screens skewed younger than that of WT controls. We acquired two additional WT controls so that the age distributions of FUS-ALS and WT patient cell lines were better matched (supplementary Table 1). With this expanded library, we screened 13 WT and 12 FUS-ALS lines using the EEA1/FUS marker set. Prediction scores highlighted strong separation between FUS-ALS and WT cell lines (AUROC = 0.83, 0.84, 0.76 for cell line, well replicate or single cell aggregation, respectively; Fig. 2a-c). Scores were consistent within cell line (18 well replicates, over 3 plates) and FUS-ALS phenotype scores ranged along a spectrum, from near WT to strongly separated (Fig. 2b). Interestingly, two FUS-ALS fibroblast cell lines (F6 and F12) were biopsied pre-symptomatically, yet were among the most strongly separated cell lines. Topranked features in classification models all captured intensity ratios between EEA1 and FUS, involving EEA1 intensity across all cellular regions (cytoplasm, DNA, full cell) and FUS intensity localized to the cytoplasm (Fig. 2d). These identified features guided a projection into twodimensional coordinate space—one axis for each marker—that highlighted distributional differences between the FUS-ALS and WT populations (Fig. 2e), consistent with the IF images (Fig. 2f). Taken together, these results demonstrate a robust separation between WT and FUS-ALS patients based on a small number of intuitive and visible cellular phenotypes.

**Figure 2:**
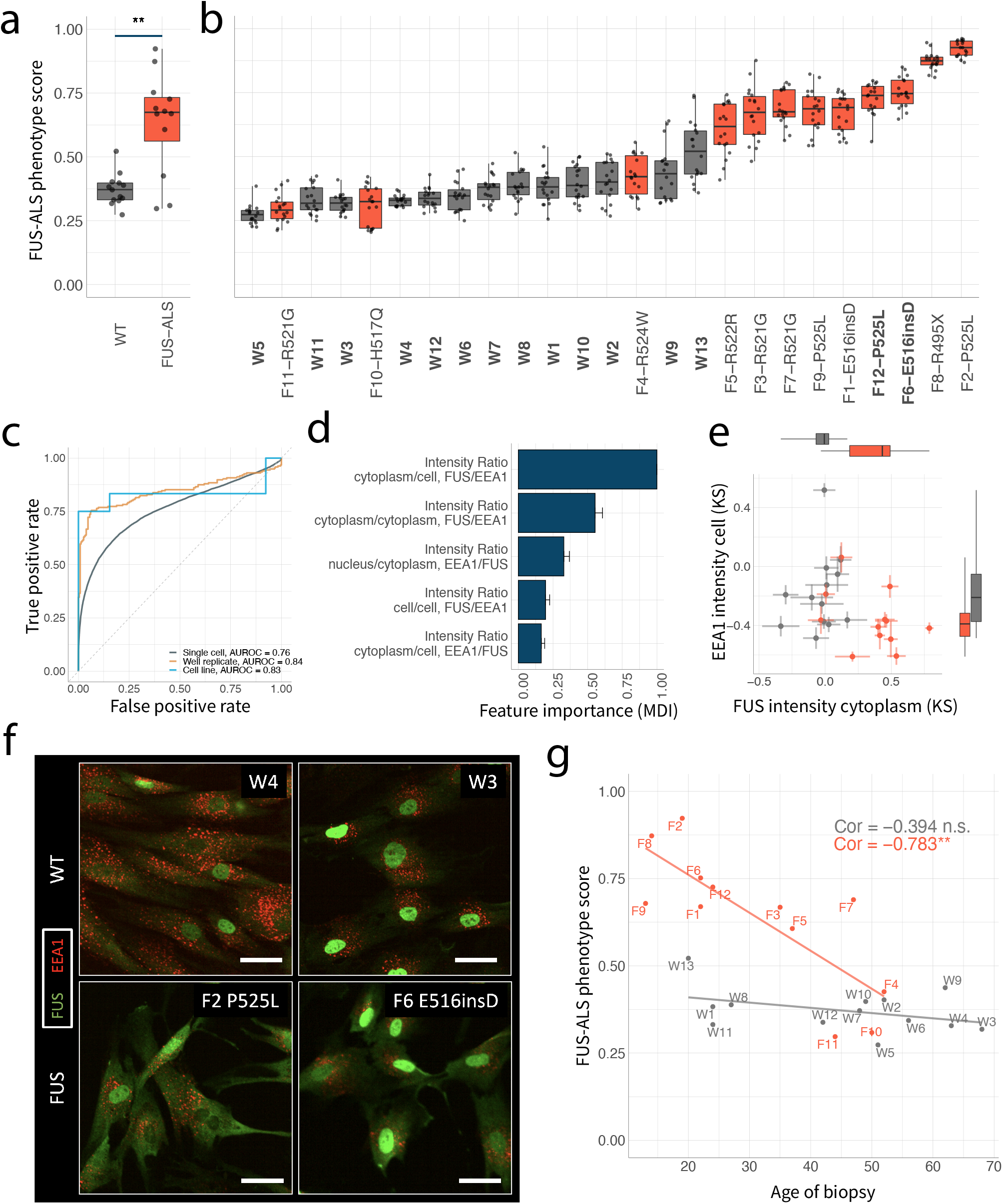
FUS-ALS separation from control WT. **a-b,** FUS-ALS phenotypic score averaged over cell line (a) or well-level replicates (b). Dots: (a) cell lines or (b) well replicates. p-value: one-sided Wilcoxon rank-sum test; ** < 0.01. **c,** ROC curve showing single cell, well or cell line classification accuracy based on FUS-ALS phenotypic scores. **d,** Feature importance averaged across all cell line-holdout models and normalized to the top performing feature. MDI: mean decrease in impurity. Error bars: 1 standard deviation across cell line-holdout models. **e,** Two-dimensional projection of FUS-ALS subtype and WT profiles. Dots: averaged Kolmogorov-Smirnov (KS) profiles over well replicates; KS relative to a WT cell line (Methods); lines: 1 standard deviation of KS profiles. **f,** Representative images of cellular phenotypes. Scale-bar: 20um. **g,** Scatterplot of age of biopsy vs. FUS-ALS phenotypic scores. p-value: one-sided t-test, Bonferroni corrected; ** < 0.01, ns > 0.05. (a-b, e, g) Colors: gray (WT), red (FUS-ALS). (a-b, e) Box plots: median (center line), interquartile range (box) and data range (whiskers).

Using our FUS-ALS classification model, we investigated how FUS-ALS phenotype scores relate to FUS-ALS pathobiology. First, we tested whether the spectrum of FUS-ALS phenotypic scores from the 25 selected patient cell lines (13 WT, 12 FUS-ALS) reflected clinical age of onset, a measure of disease severity. We compared scores against patient age at biopsy (for FUS-ALS patients, collection typically occurred within a year of disease onset; Methods). WT patient age and model scores showed no significant correlation (Pearson correlation = −0.394; one-sided t-test, Bonferroni corrected p-value = 0.275; Fig. 2g). However, a significant negative correlation was observed between FUS-ALS patient age and model prediction scores (Pearson correlation = −0.783; one-sided t-test, Bonferroni corrected p-value = 0.004; Fig. 2g). Thus, FUS-ALS phenotypic scores provide a proxy for FUS-ALS severity across a range of patient genetics: late onset FUS-ALS more closely resembles WT (Fig. 2g). These results were consistent when replacing FUS-ALS patient age of biopsy with age of diagnosis (Extended Data Fig. 3).

We next investigated whether imaging-based separation is reflected in transcriptional-based readouts. At the time image assays were performed, we additionally collected mRNA from cell lines under the same culture conditions (Methods). After sequencing, we ran differential gene expression analysis comparing WT and FUS-ALS cell lines (Methods). While EEA1 mRNA was upregulated in WT cell lines relative to FUS, consistent with our imaging-based profiles, the difference was not statistically significant (B-H FDR corrected p-value = 0.829, Fig. 3a). Similarly, CD63 and FUS mRNA expression was not significantly different between WT and FUS-ALS cell lines (B-H FDR corrected p-values = 0.997, Fig 3a). Broadening our search beyond FUS, EEA1, and CD63 highlighted only four genes that were differentially expressed in fibroblasts (B-H FDR corrected p-value < 0.05), though all with some previous association to ALS ^33–36^ (Fig. 3b). To directly evaluate the degree to which transcriptional signals generalized to new cell lines—an important requirement of our imaging platform—we trained a FUS-ALS vs. WT sparse classifier on eigengenes (Methods). The eigengene classifier achieved notably lower accuracy compared with our imaging-based model (eigengene vs. imaging classifier recall: 2/12 vs. 9/12; though, several of the cell lines accurately classified by imaging were close to the sequencing decision boundary; Fig. 3c). There was little correlation between the prediction scores from the two modalities (Fig. 3c). Thus, imaging-based separation was not reflected in transcription-based readouts and provides better separation.

**Figure 3:**
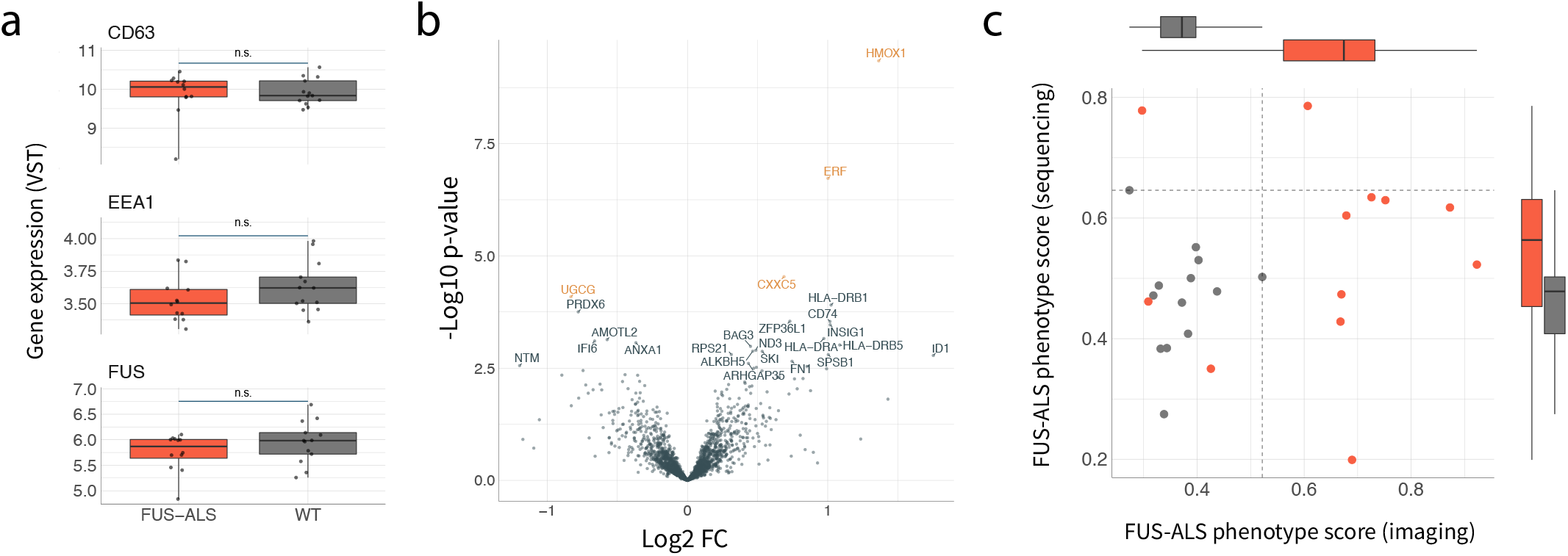
Comparison of phenotypic and transcriptional ALS profiles. **a,** Gene expression of top biomarkers (variance stabilizing transformation). Dots: individual cell lines. ns: p-value > 0.05, differential expression analysis (Deseq2; Methods). **b,** Volcano plot. X-axis: log2-fold change (FC) FUS-ALS vs WT; Y-axis: −log10 differential expression p-value. Colored genes: corrected BH corrected p-value < 0.05. **c,** Scatterplot of FUS-ALS imaging vs. RNAseq phenotypic scores. Dots: individual cell lines. Dashed lines: maximum scoring WT cell line.

Could the FUS-ALS phenotypic score offer insights into sporadic ALS patients with no known genetic cause? We had additionally screened and sequenced 11 fibroblast cell lines obtained from sporadic ALS patients with the FUS-ALS and WT cell lines above. We evaluated our FUS-ALS score on the sporadic cell lines. While the sporadic cell lines showed weaker separation from WT than FUS-ALS, five were close to the decision boundary (Fig. 4a-b). We divided sporadic cell lines into high scoring (sporadic+) and low scoring (sporadic-) groups and conducted gene set enrichment analysis^38^ for both groups along with FUS-ALS cell lines (Fig. 4b-c). Interestingly, sporadic+ cell lines showed consistent pathway enrichment with the FUS-ALS cell lines (Fig. 4c). In contrast, sporadic-showed no pathway enrichment (Fig. 4c). Thus, the FUS-ALS phenotypic score provided a way to identify sporadic subgroups that share pathway dysregulation with FUS-ALS fibroblasts.

**Figure 4:**
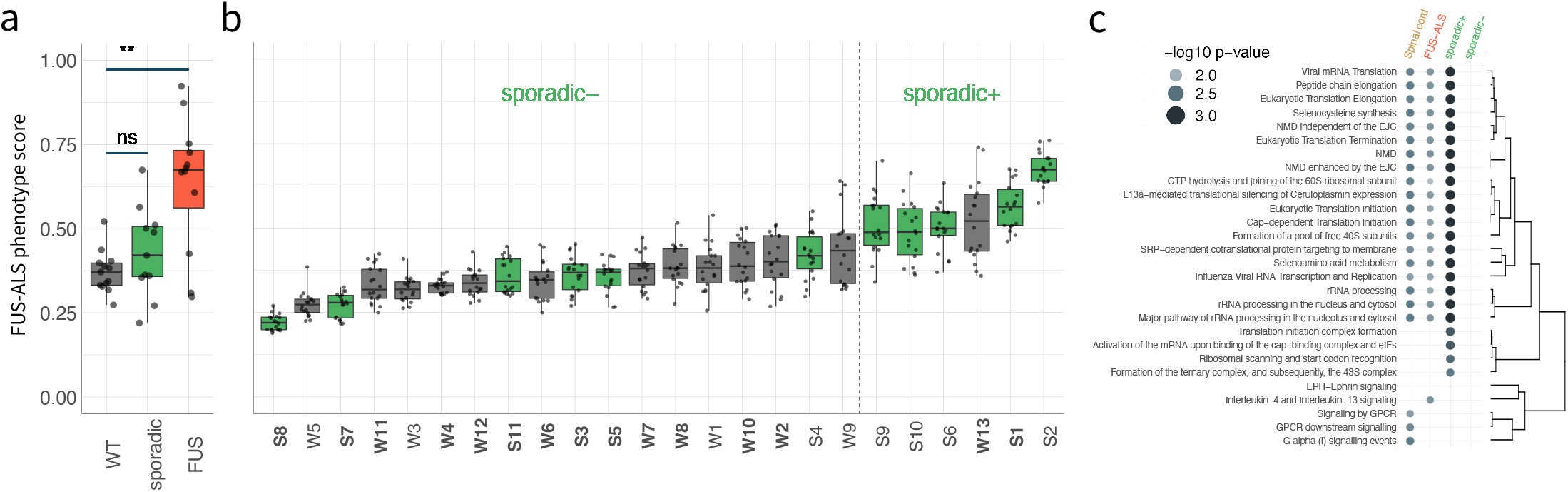
Sporadic ALS separation from control WT. **a-b,** FUS-ALS phenotypic scores averaged over cell line (a) or well-level replicates (b). Colors: gray (WT), green (sporadic ALS). Dots: (a) cell lines or (b) well replicates. p-value: one-sided Wilcoxon rank-sum test, Bonferroni corrected ** < 0.01, ns > 0.05. Box plots: median (center line), interquartile range (box) and data range (whiskers). **c,** Pathway (Reactome) enrichment by sample type. NMD: nonsense-mediated decay, EJC: exon junciton complex. Sporadic+: sporadic cell lines with top 5 FUS-ALS phenotype scores. Sporadic-: sporadic cell lines with 6 lowest FUS-ALS phenotype scores. Points size/color: −log10 BH-corrected p-value (gene set enrichment analysis, Methods). Dendrogram: hierarchical clustering using Jaccard distance defined over gene sets associated with each pathway.

To assess the relevance of fibroblast lines to ALS—a motor neuron disease—we compared our fibroblast gene expression data with a library of human spinal cord samples including various ALS genetics (Methods). While these disparate tissue types might share few changes at the level of specific genes, we reasoned that they may share dysregulation at the level of pathways. Differential expression p-values showed little correlation between spinal cord and fibroblast transcriptional data (Extended Data Fig. 4). However, gene set enrichment analysis^38^ highlighted pathways that were consistent between fibroblasts and spinal cords (19 pathways enriched in all of FUS-ALS, sporadic+, spinal cord; 28 pathways enriched in one of FUS-ALS, sporadic+, spinal cord; Methods). For example, pathways common to fibroblasts and spinal cord included nonsense-mediated decay and RNA metabolism, which have been previously implicated in ALS^39^ (Fig. 4c). Thus, both the patient-derived FUS-ALS and sporadic+ fibroblasts shared pathway dysregulation with patient spinal cord samples.

Finally, we tested the ability of our imaging-based platform to evaluate potential therapeutic agents. Specifically, we used the FUS-ALS phenotypic score to examine the effects of an antisense oligonucleotide (ASO) recently developed to treat a FUS-ALS patient^37^. (The ASO targets the sixth intron of the FUS gene and knocks down both mutant and WT FUS mRNA, Methods.) We selected the five youngest FUS-ALS patient cell lines, as they showed the greatest separation in our platform, and age matched them to the five youngest WT cell lines. Cell lines were electroporated with both a low (1uM) and a high (10uM) dose of ASO. Cells recovered for 48h before being stained and imaged with the optimal imaging marker set (Methods). qPCR analysis of the FUS mRNA confirmed a dose dependent knock-down of FUS across all cell lines (Extended Data Fig. 5a). Image analysis further confirmed that average FUS intensity per cell in the nucleus decreased (Fig. 5a). *A priori*, we had expected that the ratio of nuclear to cytoplasmic intensities would remain relatively constant with respect to the ASO treatment doses, reflecting a steady state between the two compartments. Interestingly, this was not the case—in fact, average cytoplasmic FUS levels per cell dropped slightly at low ASO dose and remained constant from low to high dose (Fig. 5a).

**Figure 5:**
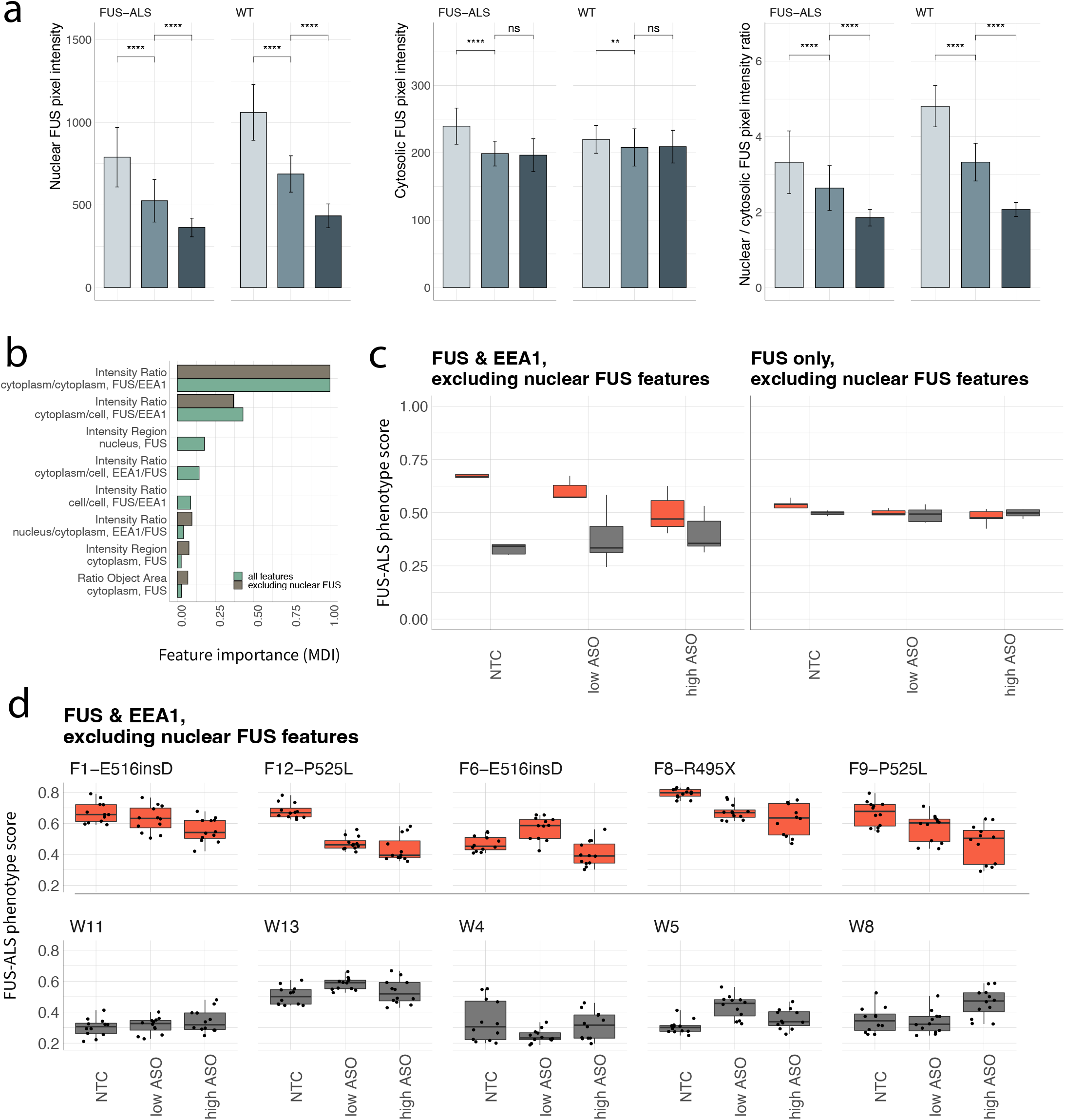
ASO treatment modulates FUS-ALS vs. WT phenotypic separation. **a,** Effect on average FUS intensity in different cellular compartments due to ASO FUS-knockdown. Bars: average intensity over group; whiskers: standard deviation across well replicates. p-value: one-sided Wilcoxon rank-sum test; ns > 0.05, ** < 0.01, **** < 0.0001. Shading: light grey (no-targeting control), medium gray (low ASO treatment, 1um), dark gray (high ASO treatment, 10um). **b,** Feature importance normalized to the top performing feature. Color: green, models trained using all features; brown, models trained with nuclear FUS features excluded. MDI: mean decrease in impurity. **c,** FUS-ALS phenotypic scores averaged within cell lines and treatments. Models trained on EP cells treated with H2O only. Left/right: models trained with/without nuclear features of FUS. **d,** FUS-ALS phenotypic scores averaged within well replicates (dots) by treatment using models without nuclear FUS features. (c,d) Colors: gray (WT), red (FUS-ALS). (c,d) Box plots: median (center line), interquartile range (box) and data range (whiskers).

Many FUS mutations disrupt the nuclear localization sequence^32^ and have been reported to reduce nuclear FUS levels ^31^. We reasoned that the large differences in nuclear FUS levels between FUS-ALS and WT (Fig. 5a) would be heavily weighted in classification models. As expected, when we fit a model using all features (nuclear, cytoplasmic, and entire cell regions), nuclear FUS levels were highly ranked with respect to feature importance (Fig. 5b). The full feature model showed WT cell lines “moving” towards FUS-ALS (Extended Data Fig. 5b). This reflected the obvious effect of ASO treatment—that knockdown of FUS mRNA leads to reduced nuclear FUS levels.

To remove the strong effect of the ASO on nuclear FUS, we fit a model excluding nuclear FUS features (i.e., restricted to cytosolic FUS and all EEA1 features). The *excluding nuclear FUS* model showed FUS-ALS cell lines “moving” towards WT (Fig. 5c). That is, the overall lowering of FUS levels through the ASO led to a more WT-like cytoplasmic phenotype in mutant cells (as measured by cytosolic EEA1 and FUS intensities; Fig. 5c-d). We noted that the separation and reversion obtained from using both EEA1 and cytosolic FUS features was considerably more pronounced than using cytosolic FUS features alone (Fig. 4c). (Motivated by these results, we re-analyzed our original FUS-ALS screen (Fig. 2) excluding nuclear features and found that the separation remained (Extended Data Fig. 6); that is, despite reading out different aspects of FUS biology, both whole cell and cytosolic FUS-only features were able to separate WT from FUS-ALS). Taken together, these results demonstrated that the platform can capture the effects of therapeutic candidates across distinct patient cell lines.

## DISCUSSION

Our profiling platform provides an approach to move from genetic to phenotypic subtyping in ALS using primary patient-derived fibroblast cell lines. We used a combined high-content imaging and machine learning platform to search over disease-relevant imaging markers for interpretable, cellular phenotypes that distinguish ALS genetic subtypes from WT. We identified EEA1 and FUS as imaging markers that separate FUS-ALS from WT (Fig. 1). We confirmed that our separation model generalizes to new WT and FUS-ALS cell lines.

The separation between FUS-ALS from WT allowed us to define an FUS-ALS phenotype score. This score measures a “distance” of each cell line to WT, quantifying a disease spectrum within FUS-ALS. Our study shows how the FUS-ALS phenotype score can be applied to reveal meaningful information about the ALS patient cell lines. First, the cell line scores were negatively correlated with age of FUS-ALS diagnosis: earlier onset patients tended to fall further from WT (Fig 2). Second, the score identified subsets of sporadic cell lines that have consistent pathway dysregulation with FUS-ALS (Fig. 4) and performed better than transcription-based scores at separating FUS-ALS vs. WT cell lines (Fig. 3). Finally, the score revealed ASO treatment effects (Fig. 5). Taken together, the FUS-ALS phenotype score provides a new lens through which to investigate ALS patient heterogeneity.

The FUS-ALS phenotypic score suggests a quantitative approach to stratify sporadic ALS patient cell lines. Sporadic ALS patients make up ~90% of reported cases. Most of these patients have no known genetic cause and exhibit clinical heterogeneity (e.g., age/location of onset and disease progression) that is difficult to study. In particular, the score groups the sporadic cell lines into subsets that are phenotypically similar or dissimilar with FUS-ALS. Interestingly, the phenotypically similar group shares pathway-level dysregulation with FUS-ALS cell lines as well as spinal cord tissue samples.

Observing how cellular perturbations affect the FUS-ALS phenotypic scores can be used to evaluate therapeutic candidates and study disease mechanisms. Fascinatingly, different modeling strategies based on the EEA1 and FUS imaging markers revealed that ASO knockdown both “pushed” WT cells towards FUS-ALS (Extended data Fig. 5b) and “reverted” FUS-ALS towards WT (Fig. 5c). The push from WT to FUS-ALS depended on reduced nuclear FUS levels, consistent with mislocalization of mutant FUS to the cytosol due to a loss of nuclear localization signal^32^. In contrast, the reversion of FUS-ALS towards WT depended on reduced cytosolic FUS levels, consistent with observed patient cases with higher cytosolic levels^31^. Decreased nuclear and increased cytosolic FUS levels in FUS-ALS have led to hypotheses of loss- and/or toxic gain-of-function (respectively) for FUS^29,38,39^. While it is not the goal of our current study to answer mechanistic questions around loss-vs. gain-of-function, the distinctions that the models captured will provide the flexibility to evaluate disease-causing phenotypes.

Studies of ALS mechanisms and/or treatment will require disease-relevant cell models that are scalable. Fibroblasts are readily obtainable from living donors, often available through biobanks, and easy to assay relative to other cell models (such as iPSCs or differentiated neurons). This allowed us to investigate a broad range of patients, representing a range of mutations and clinical variability. Our study shows that primary patient-derived fibroblasts contain disease-relevant information and will complement studies focused on genetically edited isogenic cell line pairs^40^. A corollary of our work is the ability to guide rational construction of patient cell line resources around specific phenotypic disease subtypes.

There are several limitations to our study. First, our results are based on a relatively small number of patient samples. It has been widely remarked in studies of ALS that low prevalence of the disease is a major hurdle to acquiring patient samples. Nevertheless, we endeavored to consider a diverse patient population (i.e., ALS genetics, mutation, patient age, sex) relative to available samples and test the generalization of results by adding new cell lines as they became available. The robustness of our platform and results will continue to be stress-tested as more samples become available. Second, fibroblasts are unarguably a simplified model of ALS. Primary fibroblasts have the potential—by enabling high-throughput studies across diverse patient populations—to accelerate early-stage identification of potential therapeutics. We demonstrated the relevance of fibroblasts to ALS, a motor neuron disease, and showed that pathway dysregulation is consistent with patient-derived spinal cord samples. However, as with all experimental models (including iPSCs or mouse models), future studies are needed to determine to what extent reversion of disease phenotypes to health in fibroblasts will translate to therapeutic solutions for ALS in humans. Third, our main results focus on FUS-ALS, which is a rare subpopulation (~3-6% in familial and <1% in sporadic ^4^) of ALS. However, we found preliminary evidence that other genetics, particularly a subset of C9orf72 patients, separated from WT. Extending our platform to other ALS genetic subtypes will further characterize the ALS disease landscape and provide starting points for rational selection of patient subpopulations for targeted therapies.

## Acknowledgements

We gratefully thank the ALS patients and research subjects who contributed samples, and the staff of the Columbia Motor Neuron Center, The Harvard Stem Cell Institute, and the Memory and Aging center at UCSF for banking and providing patient-derived fibroblasts. We thank the Target ALS Human Postmortem Tissue Core, New York Genome Center for Genomics of Neurodegenerative Disease, Amyotrophic Lateral Sclerosis Association and TOW Foundation for spinal cord tissue samples. We thank Valerie, Meredith and Jenifer Estess, Erin Fleming, Emily Lowry, Margot Shanahan at ProjectALS and Benjamin Hoover and Barbara Corneo at Columbia for their support of this work. We are grateful to Tom Maniatis, Spiros Liras, Kim Huard, Aimee Kao, Bill Seeley, and Altschuler and Wu lab members for helpful discussions.

H.P. and all NYGC ALS Consortium activities are supported by the ALS Association (ALSA, 19-SI-459) and the Tow Foundation. K.K. was supported the ProjectALS Research Fellowship for Innovation and Drug Discovery, M.A.R. was supported by the ProjectALS Award for Research Excellence; S.J.A. and L.F.W gratefully acknowledge support from ProjectALS, and the DOD Amyotrophic Lateral Sclerosis Research Program Therapeutic Idea Award (ALS200175).

## Competing Interests

L.F.W., M.P.J., S.J.A. are founders and SAB members of Nine Square Therapeutics. N.S. received research support from Ionis Pharmaceuticals and is a PI on the ION363 trial.

## MATERIALS AND METHODS

### Skin fibroblast extraction and cell line generation

Fibroblasts were collected under IRB-approved protocols at Columbia, Massachusetts General Hospital following protocols below.

#### Tissue collection at UCSF

Dermal tissue was provided by the University of California, San Francisco (UCSF), Memory and Aging Center, where participants enrolled in observational research studies underwent skin biopsy after providing written informed consent. Approximately two-millimeter dermal punches were performed at the inner thigh. The tissue was immersed in complete media composed of Dulbecco’s Modified Eagle Medium (DMEM) high glucose with sodium pyruvate, 10% heat inactivated fetal bovine serum (FBS), and 1% Penicillin-Streptomycin.

#### Fibroblast cell line creation at UCSF

The tissue was divided into smaller sections, approximately one to two millimeters, with surgical tools and transferred into a six-well cell culture plate. A glass coverslip was gently placed over the tissue and complete DMEM media was added into the wells. Cultures were incubated in a humidified chamber at 37°C and 5% CO2, left undisturbed for the first week, then fed every two to three days with complete DMEM media. After approximately three weeks in culture, the cells were collected, expanded, and banked in liquid nitrogen with complete DMEM media and 10% DMSO.

#### IRB Info for UCSF

Dermal biopsies were obtained from subjects with written informed consent and the research was approved by the University of California, San Francisco Committee on Human Research (#10-00234).

#### Funding at UCSF

Bruce Miller (UCSF Memory and Aging Center, NIH/NIA P50 AG023501); Robert Farese, Fen-Biao Gao, Anna Karydas, and Sandra Almeida (California Institute for Regenerative Medicine); Yadong Huang (J. David Gladstone Institutes)

#### Tissue collection at Columbia

Dermal tissue was provided by Columbia University, from the Biorepository for the Study of Neuromuscular Disorders where participants enrolled in observational research studies underwent skin biopsy after providing written informed consent. The standard biopsy size is 3mm and is obtained with a punch. Typically one punch biopsy, is obtained. The skin biopsy sample is immediately immersed in tissue culture media and transported to the tissue culture lab. The fibroblast culture is established by adhering the biopsy to the bottom of a tissue culture plate in tissue culture media with a sterile needle. Established fibroblast cultures are then stored in cryopreservative in liquid nitrogen.

#### IRB Info for Columbia University

Dermal biopsies were obtained from subjects with written informed consent and the research was approved by Columbia University IRB# AAAK2000.

#### Tissue collection at MGH

TBD.

### Spinal cord data collection

Spinal cord samples were provided through the New York Genome Center ALS consortium. In total, we considered 202 (41 WT control, 161 ALS) cervical spinal cord samples diverse with respect to ALS mutations, age distribution, and sex (supplementary Table 4).

### Cell line culturing

Cell lines received from various institutions (UCSF, Columbia, MGH) were thawed, and expanded for future use. This established low passage replicate vials for our experiments.

All fibroblast cell lines were cultured in DMEM/F-12 (Thermo Fisher Scientific, 11320-033) supplemented with 20% FBS (Gemini, Cat# COAA33H00I), 1% GlutaMAX (2 mM) and 1% Pen-Strep at 37deg C, 5% CO2 and 80% humidity. Cell lines were thawed into 6-well plates at around 3e5 cells/mL, media was exchanged after 1 day, and cells were passaged when 100% confluent. Passaging was done at a minimum of every 3 days or at a maximum of every 7 days. Cells were only cultured up to a maximum of 20 passages. For each cell line, a new vial of cells was thawed per each batch of experiment.

### General screening setup

For each screen, all cell lines were thawed 10-14 days before the experiment and passaged when confluent. To minimize variability, all lines were split at the same time, 3-5 days before plating, at 5-7e5 cells/mL. At the start of each screen, cell lines were split and seeded into 384-well plates at 3.5e5 cells/mL. Cells were then grown for 24h and fixed for subsequent staining.

All screens used randomized location of cell lines on the plate. Plate replicate layouts were consistent within a screen and varied between screens to confirm that there is no plate position effect. Experiments were repeated (data not shown) with different plate maps to confirm that there is no plate effect. Each plate consisted of more than 4 replicate wells for each cell line. No positional effects were observed. No edge wells were used. To establish the robustness of our screens, experiments were also repeated by another person and using different microscopes.

#### ASO treatment

As above, cell lines used in screens were thawed 10-14 days before the experiment and passaged regularly. To minimize variability, all lines were split at the same time 3-5 days before the experiment at 5-7e5 cells/mL. For ASO treatment, cells were split and electroporated at 2.5e5 cell/mL concentration using a Bio-Rad Gene Pulser Xcell Electroporation System at 160V for 6ms using 0.2cm cuvettes. Cell lines were then seeded into 384-well plates at 3.5e5 cells/mL or 4.5e5 cells/mL for untreated or treated wells, respectively. Each plate consisted of more than 4 replicate wells for each cell line and condition. Cells were then grown for 48h and fixed for subsequent staining. Additionally, 12-well (6e5 cells/mL) for Western Blot analysis and 24-well (3e5 cells/ml) for qPCR analysis.

### Immunofluorescence protocol

All solutions for IF staining were made in 1X Dulbecco’s PBS (DPBS) with added Mg and Ca (10X PBS diluted with ddH20, Gibco 14-080-055). All wash steps were performed using 1X DPBS. Primary and directly conjugated antibodies were purchased as outlined in Table 2 and stored according to supplied data sheets. Cells were incubated with 200nM Mitotracker for 30 min before fixing them in 4% paraformaldehyde (Electron Microscopy Sciences Cat# 15710, Hatfield, PA) for 15 min, washed and permeabilized with 0.2% Triton at room temperature for 15 min and washed twice. Cells were blocked for 1h at room temperature with Blocking buffer made up of 0.1% Triton and 3% BSA. Primary antibodies (supplementary Table 2) were diluted in Antibody Buffer made up of 0.1% Triton and 1% BSA and incubated overnight at 4 °C. Samples were then washed and incubated for 1 h at room temperature with secondary antibodies (supplementary Table 2) and Hoechst and then washed again. Cells were then kept in PBS until imaged.

### Imaging

Plates were imaged on a Nikon Ti inverted epifluorescence microscope using plate-adjusted parameters. Images were captured DAPI, FITC, TRITC, and Cy5 channels.

### qPCR

Following protocol for iScript Reverse Transcription Supermix for RT-qPCR and using SsoAdvanced Universal SYBR Green Supermix.

### Sequencing

mRNA sequencing was completed by Genewiz/Azenta.

### Genewiz/Azenta (RNAseq)

Sample QC, library preparations and sequencing reactions were conducted at GENEWIZ/Azenta, LLC. (South Plainfield, NJ, USA) as follows:

#### QC, Library Preparation with PolyA Selection and HiSeq Sequencing

The RNA samples received were quantified using Qubit 2.0 Fluorometer (ThermoFisher Scientific, Waltham, MA, USA) and RNA integrity was checked using TapeStation (Agilent Technologies, Palo Alto, CA,USA). The RNA sequencing library was prepared using the NEBNext Ultra II RNA Library Prep Kit for Illumina using manufacturer’s instructions (New England Biolabs, Ipswich, MA, USA). Briefly, mRNAs were initially enriched with Oligod(T) beads. Enriched mRNAs were fragmented for 15 minutes at 94 °C. First strand and second strand cDNA were subsequently synthesized. cDNA fragments were end repaired and adenylated at 3’ends, and universal adapters were ligated to cDNA fragments, followed by index addition and library enrichment by PCR with limited cycles. The sequencing library was validated on the Agilent TapeStation (Agilent Technologies, Palo Alto, CA, USA), and quantified by using Qubit 2.0 Fluorometer (ThermoFisher Scientific, Waltham, MA, USA) as well as by quantitative PCR (KAPA Biosystems, Wilmington, MA, USA). The sequencing libraries were multiplexed and clustered onto 3 flowcell lanes. After clustering, the flowcell was loaded on the Illumina HiSeq instrument (4000 or equivalent) according to manufacturer’s instructions. The samples were sequenced using a 2×150bp Paired End (PE) configuration. Image analysis and base calling were conducted by the HiSeq Control Software (HCS). Raw sequence data (.bcl files) generated from Illumina HiSeq was converted into fastq files and de-multiplexed using Illumina’s bcl2fastq 2.20 software. One mismatch was allowed for index sequence identification.

## COMPUTATIONAL METHODS

### IMAGE ANALYSIS

#### Image processing of fluorescence microscopy images

In our iterative search we assayed 13 markers, multiplexed in sets of 4, across three screens: two segmentation markers (Hoechst; nuclear stain and Mitotracker; cellular outline) and two “disease-relevant” markers (supplementary Table 2). We imaged 12 fields of view from each well at 20X magnification. Raw images were flat-field corrected^41^ and background subtracted using a rolling ball algorithm^42^. Individual cellular regions were segmented from corrected images using a watershed-based algorithm^43^. For each cell, we extracted 222 quantitative features describing the spatial distribution of the two disease relevant markers (supplementary Table 3). These features were selected to provide intuitive measures of ALS-relevant phenotypes (e.g., protein intensity, localization, aggregation).

Correlation among features can make it difficult to interpret how qualitative cellular attributes contribute to model predictions. Moreover, large and redundant feature sets can lead to instability in model feature selection^44^. To mitigate these issues for modeling, we grouped features into 54 related categories (e.g., intensity in DNA region, object area in non-DNA region (supplementary Table 3)) and compressed features within each category into “eigenfeatures” using PCA, taking principal components that explained 95% of the variance in the category (typically 1-3 eigenfeatures). Thus, the feature sets used in our models summarized the phenotypes of individual cells through 54 feature categories, each consisting of a varying number of eigenfeatures. Feature reduction was performed for each screen. (Classification models, described below, were trained and evaluated within each screen. Any subsequent comparisons across screens were made at the level of feature groupings and model predictions.)

For figures showing 2D projections of samples (e.g. Fig. 1d, Fig. 2e), we summarized the distribution of raw features using a non-parametric signed Kolmogorov-Smirnov (KS) statistic. For each feature, we compared the distributions (over individual cells) of a selected well and reference WT cell line on the same plate.

#### Imaging-based classifiers

##### Model choice

We assessed the strength of separation between ALS (all subtypes) vs. WT using iterative random forest (iRF) classifiers^25^ trained on the image-based features defined above. Effectively, we projected high-dimensional image-based profiles onto a 1-dimensional score measuring predicted probability ALS. Our particular model choice has the benefits of: (i) imposing sparsity constraints that encourage models to make predictions from a small set of features; (ii) providing clear measures of each feature’s contribution to model predictions, measured as the mean decrease in Gini impurity (MDI); and (iii) provide a natural hierarchical approach to compare ALS vs WT, where each of these groups can have within-group heterogeneity (e.g. across different ALS subtypes, cell lines within each subtype, cells within one cell line). Taken together, these benefits allow for expressive and interpretable models, enabling us to identify biological signatures that differentiate ALS from WT cells.

##### Scoring

Specifically, each cell is represented as a high-dimensional point in eigenfeature vector space. iRF models were trained to classify ALS vs. WT cells using the R package iRF with two iterations and the ranger random forest classifier. The remainder of iRF parameters were set to their default values. We scored cells according to their predicted probability of being ALS as estimated by iRF models. When comparing ALS subtypes with WT, scores were averaged across cells from each cell line (e.g., Fig 2a). When comparing replicates within a cell line, scores were averaged across cells from each well replicate (e.g., Fig 2b). To compare scores between ALS and WT samples, we report both area under the receiver operating curve (AUROC)—defined for single cell, well replicate, and cell line averaged predictions—as well as recall with prediction scores thresholded at the maximum WT score (i.e. specificity = 1).

For each iRF model, we evaluated feature importance using MDI. To rank feature importance across all cell line models (see generalizability below), MDI feature importances were normalized to a 0-1 scale and averaged across models.

##### Generalizability

To ensure that scores measured the generalizability of learned phenotypes to new cell lines, scores were computed using a leave-one-cell-line-out sample splitting strategy. Predictions for a given cell line were generated from an iRF fit with that cell line held-out from training. As a result, all prediction scores are reported on cell lines that were unobserved during model training. We repeated this process (i.e., training and evaluating models) for each cell line to score its probability of being ALS. This leave-one-cell-line-out strategy has the potential to introduce bias in model predictions if different training sets contain different proportions of ALS vs. WT. To ensure balanced representation of cell line replicates and ALS vs. WT classes across each instance of model training, we randomly sampled 100 cells from each well in a screen and subsequently down-sampled the resulting dataset (to match the less prevalent class) for class balance.

#### Plate level technical variability

We evaluated the degree to which biological (i.e., ALS subtype vs. WT) variability exceeds plate-plate technical variability using a leave-one-plate-out model fitting strategy within each screen (Extended Data Fig. 2). This strategy trains iRF classifiers to discriminate between ALS and WT using cells from one replicate plate and evaluates predictions on cells from a different, held-out plate. Comparing the predicted scores between ALS and WT cells in the held-out plate allowed us to assess how well learned phenotypes generalize to a repeated experiment that is identical except with respect to plate-level variation.

### SEQUENCING ANALYSIS

#### RNA-seq data processing

For both the fibroblast and spinal cord datasets, reads were aligned to UCSC reference genome hg38 human genome using STAR 2.7.10a with default parameter settings. The resulting raw read counts for each gene were normalized as transcripts per million (TPM), with gene length given as the sum of exon length. Normalized genes were subsequently filtered by expression level, retaining protein-coding genes with median (over cell line samples) log(TPM + 1) > 2.5. The expression threshold was selected to ensure that both EEA1 and FUS were maintained in the dataset and resulted in 4228 and 5447 genes maintained in the fibroblast and spinal cord datasets respectively. Differential expression analyses were performed using the R package DESeq2^45^ controlling for the site at which each sample was collected. B-H corrected FDR p-value threshold of p < 0.05 was used to determine significance. Gene expression levels were subsequently transformed for eigengene classifier analysis using the variance stabilizing transformation from DESeq2 with fitType=“local”.

#### Eigengene classifier

Due to the high ratio of genes to samples in our fibroblast gene expression dataset and resulting instability of classification models in such settings^44^, we opted for an analysis approach that reduced the data dimensionality prior to model fitting. Expression levels of individual genes were mapped to 25 “eigengenes” using PCA. Each eigengene corresponds to a weighted combination of individual genes and cell lines are represented as a 25-dimensional vector of eigengene “expression” levels.

Using the eigengene-transformed data, we trained a set of sparse logistic regression classifiers with l1 penalty on model coefficients using the R package glmnet. The penalty parameter λ was selected using 3-fold cross validation (CV). When the penalty parameter as estimated by CV resulted in a null model, we selected the largest value (i.e., most severe penalty) that resulted in a non-null model. As in the case of our imaging classifier, we computed prediction scores for a given cell line using a model trained with that cell line removed during training. To ensure consistent class balance across all models, data were up-sampled for class balance during model training.

#### Gene set enrichment analyses

Gene set enrichment analyses (GSEAs) were run using the R package fgsea to assess the enrichment of Reactome pathways^38^. Genes were ranked by −log10 unadjusted p-value from our differential expression analysis. Parameters for minimum and maximum pathway size were set to 25 and 100 respectively.

#### ASO Analysis

For the ASO analysis (Fig. 5, Extended Data Fig. 5), all models were trained on electroporated cell lines treated with H_2_O. FUS-ALS phenotype scores were subsequently evaluated for electroporated cell lines treated with non-targeting control, low ASO, and high ASO.

## Extended Data figures

**Figure E1:**
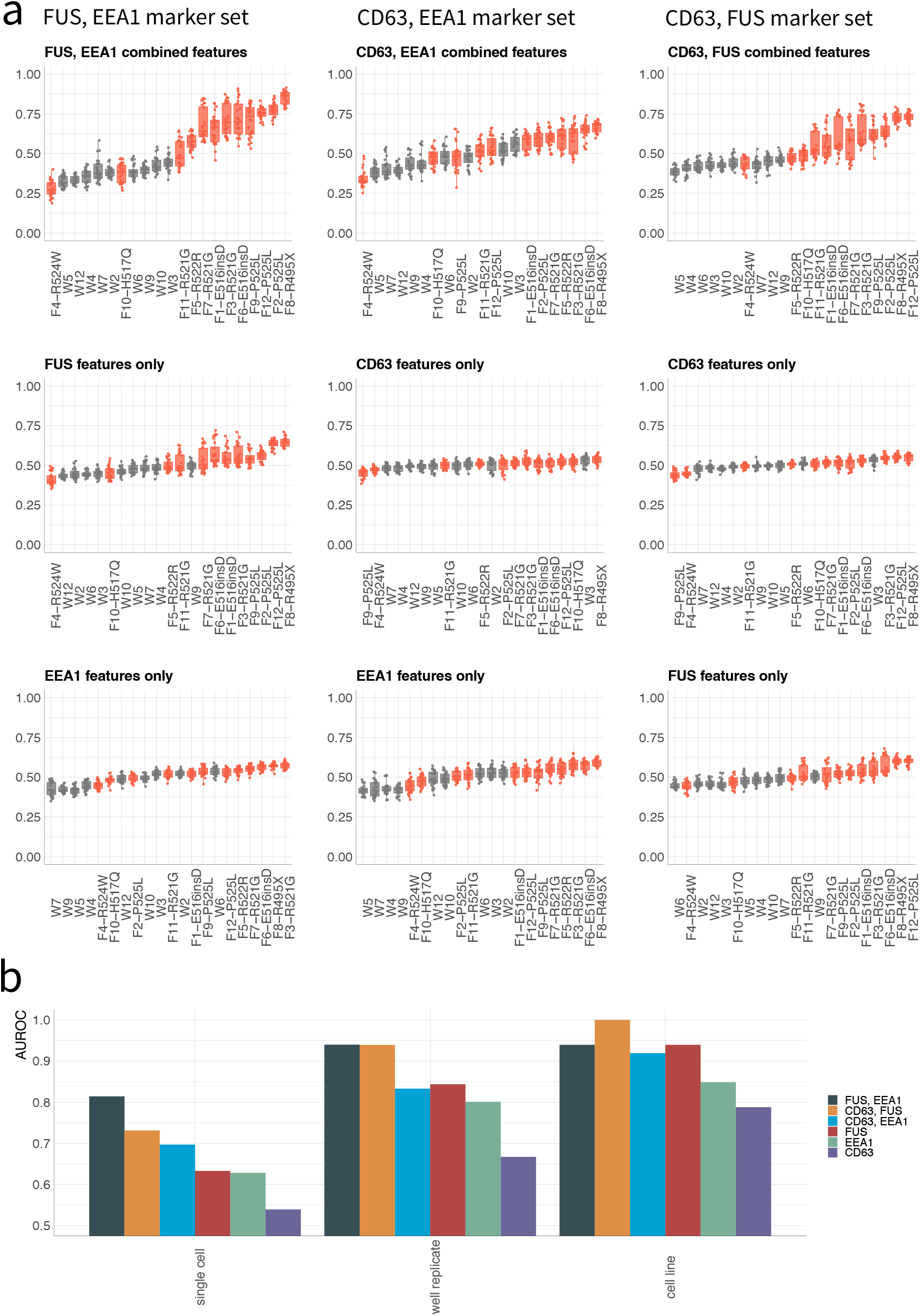
FUS-ALS separation from control WT by marker set. **a,** FUS-ALS phenotypic score for all pair-wise combinations of FUS, EEA1, and CD63 (top row) and single imaging markers (bottom two rows) averaged over well-level replicates. Colors: gray (WT), red (FUS-ALS). Dots: well replicates. Box plots: median (center line), interquartile range (box) and data range (whiskers). **b,** AUROC by marker/marker pair for single cell (left), well replicate (middle), and cell line (right) classification. F4, which is not separable using any marker/marker pair removed before computing AUROC.

**Figure E2:**
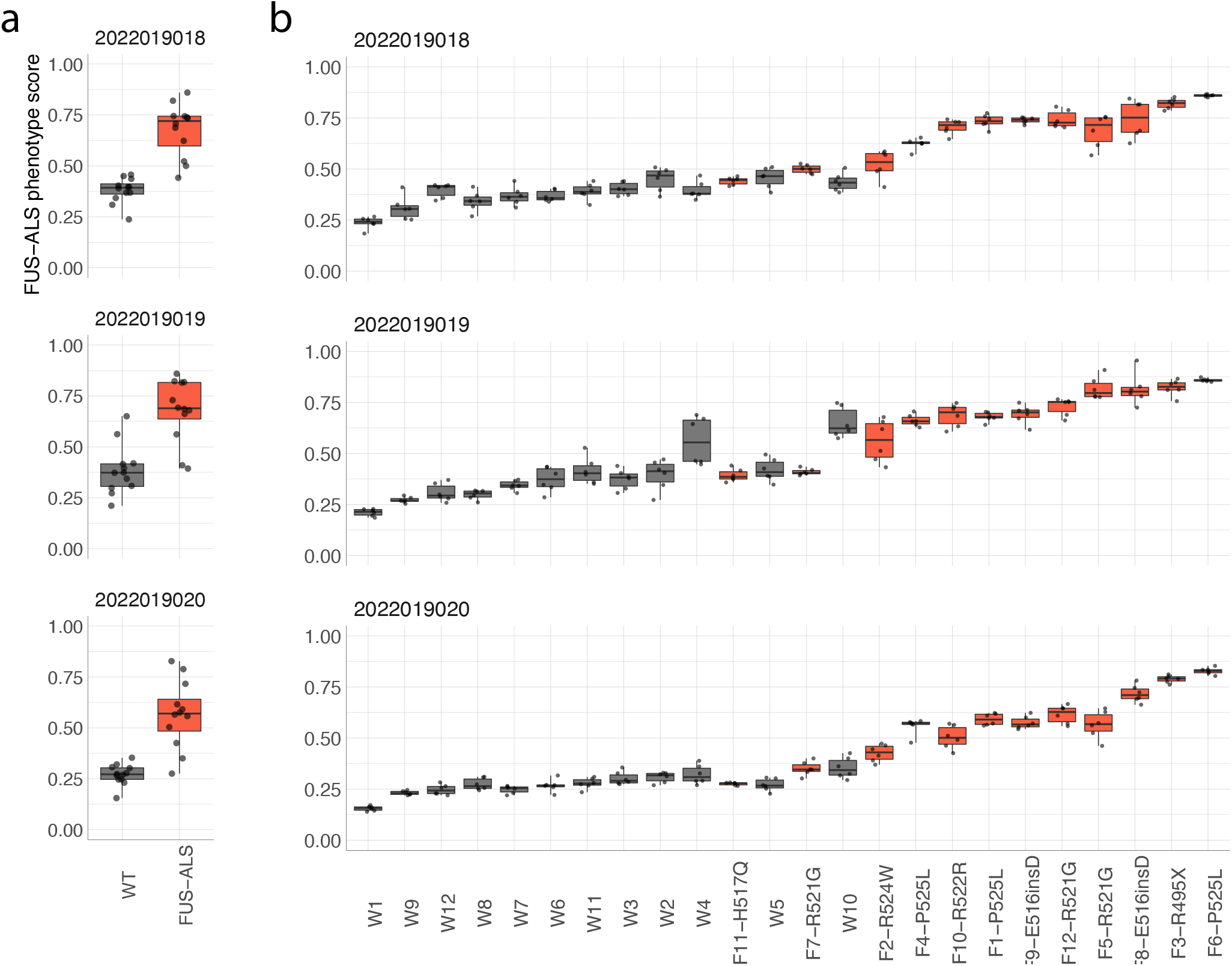
FUS-ALS phenotype scores evaluated on held-out imaging plate. **a-b,** Scores averaged over cell line (a) and well replicate (b). Dots: (a) cell lines or (b) well replicates. Box plots: median (center line), interquartile range (box) and data range (whiskers). Predictions in each imaging plate (rows) were generated using models trained on data from the remaining two plates.

**Figure E3:**
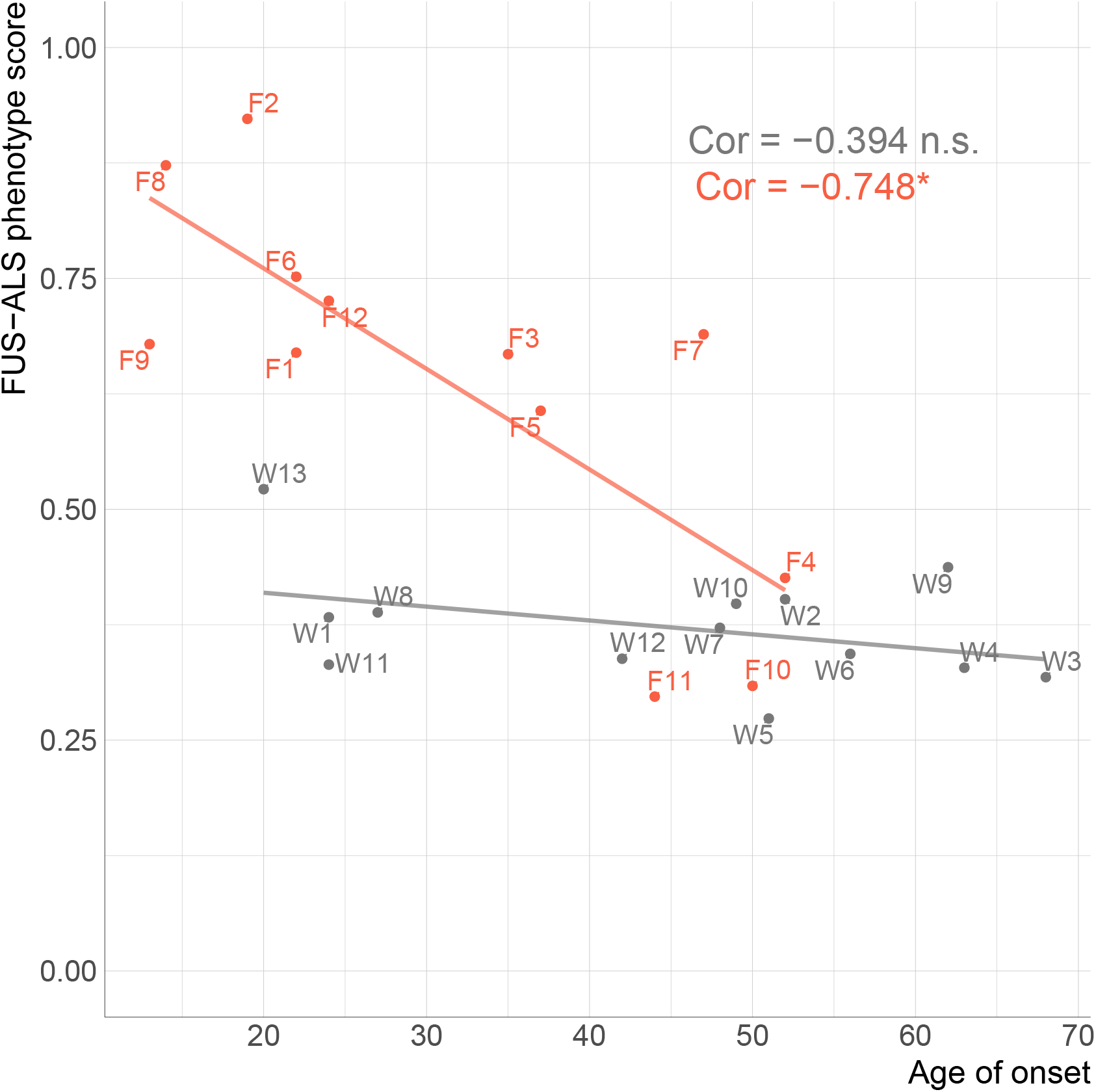
FUS-ALS phenotypic score vs. age of disease onset. FUS-ALS phenotypic score averaged over cell line vs. patient age of onset (FUS-ALS) or age of biopsy (WT). (For patient age of biopsy for FUS-ALS, compare to Fig. 2f.) Colors: gray (WT), red (FUS-ALS). p-value: one-sided t-test, Bonferroni corrected; * < 0.05. p-value FUS-ALS = 0.012, p-value WT = 0.275.

**Figure E4:**
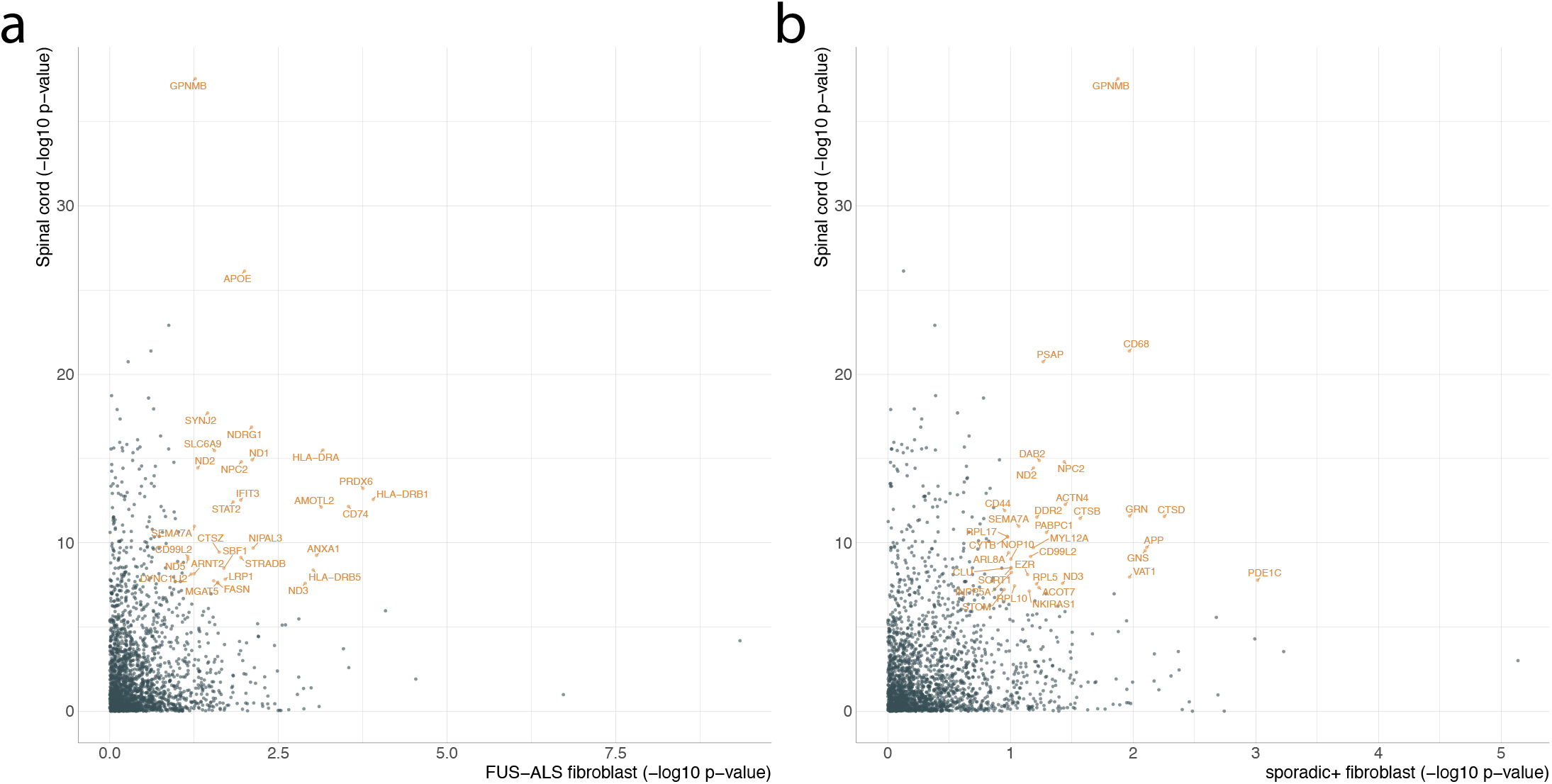
Spinal cord vs. fibroblast gene enrichment. **a-b,** Differential expression −log10 p-value in FUS-ALS fibroblasts (a) and sporadic+ fibroblasts (b) (X-axis) and spinal cord (Y-axis) samples. Highlighted genes appear in the top 10^th^ percentile of both the fibroblast and spinal cord differential expression ranked lists.

**Figure E5:**
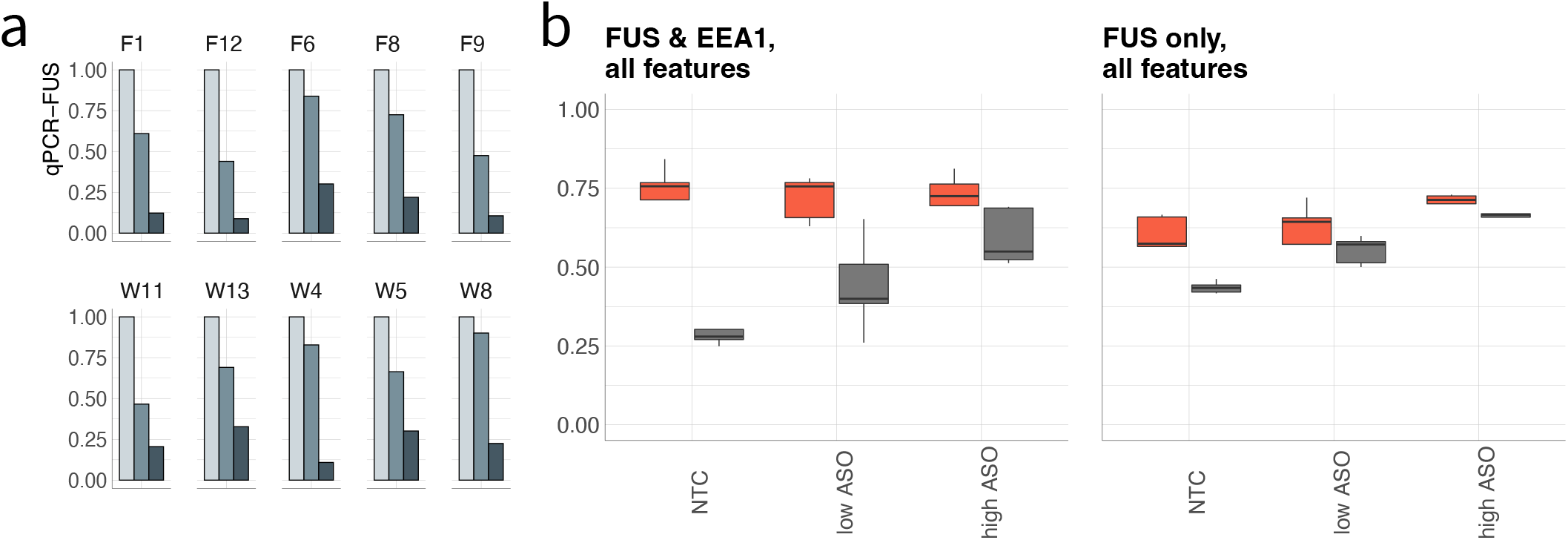
ASO treatment modulates FUS-ALS vs. WT phenotypic separation. **a,** qPCR of FUS fold-change levels after ASO FUS-knockdown. Values normalized to GAPDH and non-targeting control (Methods). EP: electroporation; NTC: non-targeting control. Shading: light grey (no-targeting control), medium gray (low ASO treatment, 1um), dark gray (high ASO treatment, 10um). **b,** FUS-ALS phenotypic scores averaged within cell lines and treatments. Models trained on EP cells treated with H2O only. Left/right: models trained with/without EEA1 features.

**Figure E6:**
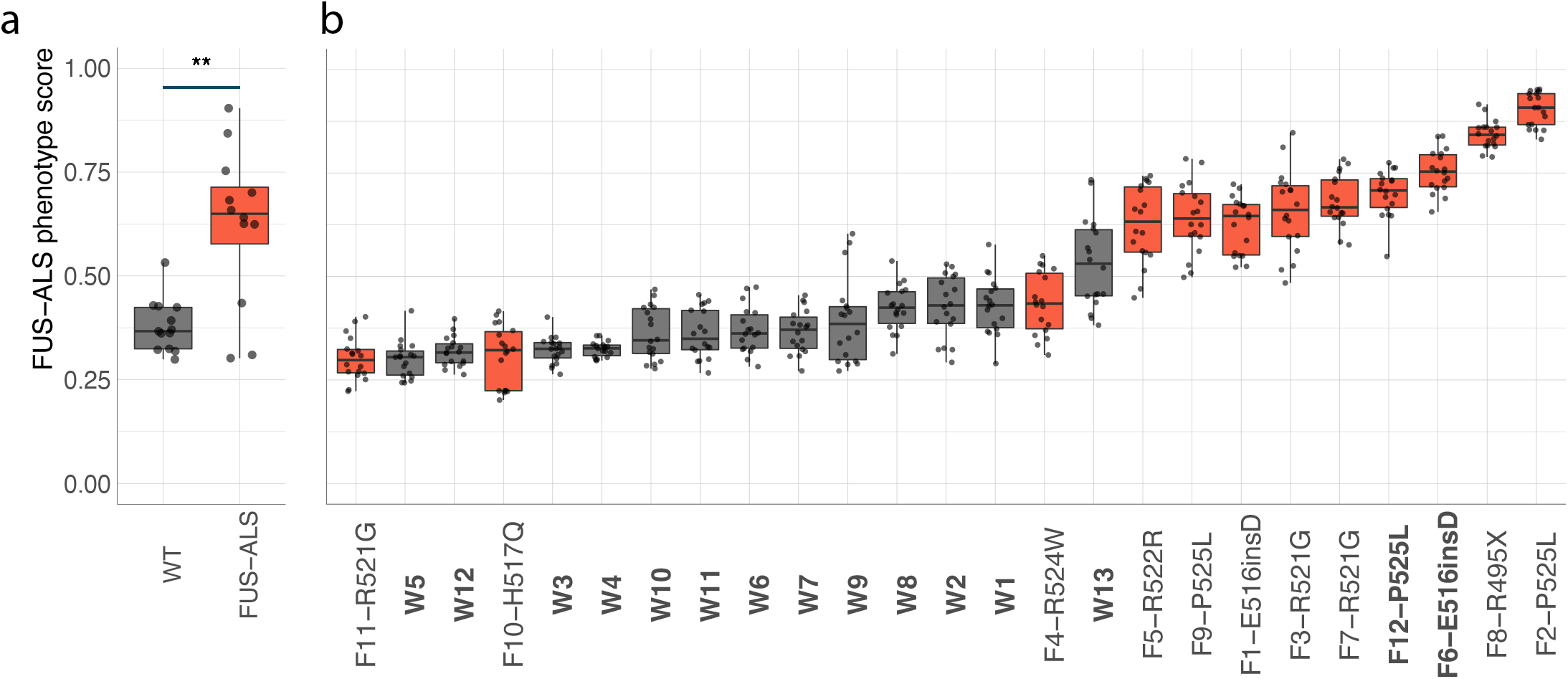
FUS-ALS separation from control WT, excluding nuclear FUS phenotypic features. **a-b,** FUS-ALS phenotypic score averaged over cell line (a) or well-level replicates (b). Models trained on data excluding FUS nuclear features. Colors: gray (WT), red (FUS-ALS). Dots: (a) cell lines or (b) well replicates. p-value: one-sided Wilcoxon rank-sum test; ** < 0.01. Box plots: median (center line), interquartile range (box) and data range (whiskers).

## Notes

### Competing Interest Statement

L.F.W., M.P.J., S.J.A. are founders and SAB members ofNine Square Therapeutics. N.S.received research support from Ionis Pharmaceuticals and is a PI on the ION363 trial.

### Summary of Updates

Author affiliations updated

## BIBLIOGRAPHY

1. Appel, S. H., Beers, D. R. & Zhao, W. Amyotrophic lateral sclerosis is a systemic disease: peripheral contributions to inflammation-mediated neurodegeneration. Curr Opin Neurol 34, 765–772 (2021).

2. Brown, R. H. & Al-Chalabi, A. Amyotrophic Lateral Sclerosis. N Engl J Med 377, 162–172 (2017).

3. Laferriere, F. & Polymenidou, M. Advances and challenges in understanding the multifaceted pathogenesis of amyotrophic lateral sclerosis. Swiss Med Wkly 145, w14054 (2015).

4. Mejzini, R. et al. ALS Genetics, Mechanisms, and Therapeutics: Where Are We Now? Front Neurosci 13, 1310–1310 (2019).

5. Weishaupt, J. H., Hyman, T. & Dikic, I. Common Molecular Pathways in Amyotrophic Lateral Sclerosis and Frontotemporal Dementia. Trends Mol Med 22, 769–783 (2016).

6. Baxi, E. G. et al. Answer ALS, a large-scale resource for sporadic and familial ALS combining clinical and multi-omics data from induced pluripotent cell lines. Nat Neurosci 25, 226–237 (2022).

7. Vance, C. et al. Mutations in FUS, an RNA Processing Protein, Cause Familial Amyotrophic Lateral Sclerosis Type 6. Science 323, 1208–1211 (2009).

8. Kwiatkowski, T. J. et al. Mutations in the FUS/TLS gene on chromosome 16 cause familial amyotrophic lateral sclerosis. Science 323, 1205–8 (2009).

9. DeJesus-Hernandez, M. et al. Expanded GGGGCC Hexanucleotide Repeat in Noncoding Region of C9ORF72 Causes Chromosome 9p-Linked FTD and ALS. Neuron 72, 245–256 (2011).

10. Freischmidt, A. et al. Haploinsufficiency of TBK1 causes familial ALS and fronto-temporal dementia. Nat Neurosci 18, 631–636 (2015).

11. Neumann, M. et al. Ubiquitinated TDP-43 in Frontotemporal Lobar Degeneration and Amyotrophic Lateral Sclerosis. Science 314, 130–133 (2006).

12. Sreedharan, J. et al. TDP-43 Mutations in Familial and Sporadic Amyotrophic Lateral Sclerosis. Science 319, 1668–1672 (2008).

13. Wang, W.-Y. et al. Interaction of FUS and HDAC1 regulates DNA damage response and repair in neurons. Nat Neurosci 16, 1383–1391 (2013).

14. Conte, A. et al. P525L FUS mutation is consistently associated with a severe form of juvenile amyotrophic lateral sclerosis. Neuromuscul Disord 22, 73–5 (2012).

15. Masrori, P. & Damme, P. V. Amyotrophic lateral sclerosis: a clinical review. Eur J Neurol 27, 1918–1929 (2020).

16. Laferriere, F. & Polymenidou, M. Advances and challenges in understanding the multifaceted pathogenesis of amyotrophic lateral sclerosis. Swiss Med Wkly 145, w14054–w14054 (2015).

17. Maniatis, S. et al. Spatiotemporal dynamics of molecular pathology in amyotrophic lateral sclerosis. Science 364, 89–93 (2019).

18. Wang, J. C., Ramaswami, G. & Geschwind, D. H. Gene co-expression network analysis in human spinal cord highlights mechanisms underlying amyotrophic lateral sclerosis susceptibility. Sci Rep-uk 11, 5748 (2021).

19. Jones, A. R. et al. Stratified gene expression analysis identifies major amyotrophic lateral sclerosis genes. Neurobiol Aging 36, 2006.e1–2006.e9 (2015).

20. Morgan, S., Duguez, S. & Duddy, W. Personalized Medicine and Molecular Interaction Networks in Amyotrophic Lateral Sclerosis (ALS): Current Knowledge. J Personalized Medicine 8, 44 (2018).

21. Tam, O. H. et al. Postmortem Cortex Samples Identify Distinct Molecular Subtypes of ALS: Retrotransposon Activation, Oxidative Stress, and Activated Glia. Cell Rep 29, 1164–1177 e5 (2019).

22. Humphrey, J. et al. FUS ALS-causative mutations impair FUS autoregulation and splicing factor networks through intron retention. Nucleic Acids Res 48, 6889–6905 (2020).

23. Schiff, L. et al. Integrating deep learning and unbiased automated high-content screening to identify complex disease signatures in human fibroblasts. Nat Commun 13, 1590 (2022).

24. Codron, P. et al. Primary fibroblasts derived from sporadic amyotrophic lateral sclerosis patients do not show ALS cytological lesions. Amyotroph Lateral Scler Frontotemporal Degener 19, 446–456 (2018).

25. Basu, S., Kumbier, K., Brown, J. B. & Yu, B. Iterative random forests to discover predictive and stable high-order interactions. Proceedings of the National Academy of Sciences 115, 1943–1948 (2018).

26. Breiman, L. Random forests. Machine learning 45, 5–32 (2001).

27. Mejzini, R. et al. ALS Genetics, Mechanisms, and Therapeutics: Where Are We Now? Front Neurosci 13, 1310 (2019).

28. Marrone, L. et al. Isogenic FUS-eGFP iPSC Reporter Lines Enable Quantification of FUS Stress Granule Pathology that Is Rescued by Drugs Inducing Autophagy. Stem Cell Reports 10, 375–389 (2018).

29. Lin, Y. C. et al. Interactions between ALS-linked FUS and nucleoporins are associated with defects in the nucleocytoplasmic transport pathway. Nat Neurosci 24, 1077–1088 (2021).

30. Liu, M. L., Zang, T. & Zhang, C. L. Direct Lineage Reprogramming Reveals Disease-Specific Phenotypes of Motor Neurons from Human ALS Patients. Cell Rep 14, 115–128 (2016).

31. Mackenzie, I. R. et al. Pathological heterogeneity in amyotrophic lateral sclerosis with FUS mutations: two distinct patterns correlating with disease severity and mutation. Acta Neuropathol 122, 87–98 (2011).

32. Tyzack, G. E. et al. Widespread FUS mislocalization is a molecular hallmark of amyotrophic lateral sclerosis. Brain 142, 2572–2580 (2019).

33. Lee, J., Kannagi, M., Ferrante, R. J., Kowall, N. W. & Ryu, H. Activation of Ets-2 by oxidative stress induces Bcl-xL expression and accounts for glial survival in amyotrophic lateral sclerosis. Faseb J Official Publ Fed Am Soc Exp Biology 23, 1739–49 (2009).

34. Bouscary, A. et al. Ambroxol Hydrochloride Improves Motor Functions and Extends Survival in a Mouse Model of Familial Amyotrophic Lateral Sclerosis. Front Pharmacol 10, 883 (2019).

35. Jimenez-Villegas, J. et al. NRF2 as a therapeutic opportunity to impact in the molecular roadmap of ALS. Free Radic Biol Med 173, 125–141 (2021).

36. Nabais, M. F. et al. Significant out-of-sample classification from methylation profile scoring for amyotrophic lateral sclerosis. Npj Genom Medicine 5, 10 (2020).

37. Korobeynikov, V. A., Lyashchenko, A. K., Blanco-Redondo, B., Jafar-Nejad, P. & Shneider, N. A. Antisense oligonucleotide silencing of FUS expression as a therapeutic approach in amyotrophic lateral sclerosis. Nat Med 28, 104–116 (2022).

38. Sanjuan-Ruiz, I. et al. Wild-type FUS corrects ALS-like disease induced by cytoplasmic mutant FUS through autoregulation. Mol Neurodegener 16, 61 (2021).

39. Ward, C. L. et al. A loss of FUS/TLS function leads to impaired cellular proliferation. Cell Death Dis 5, e1572 (2014).

40. Baxi, E. G. et al. Answer ALS, a large-scale resource for sporadic and familial ALS combining clinical and multi-omics data from induced pluripotent cell lines. Nat Neurosci 25, 226–237 (2022).

41. Zwier, J., Rooij, G. V., Hofstraat, J. & Brakenhoff, G. Image calibration in fluorescence microscopy. Journal of Microscopy 216, 15–24 (2004).

42. Sternberg, S. R. Biomedical image processing. Computer 16, 22–34 (1983).

43. Vincent, L. & Soille, P. Watersheds in digital spaces: an efficient algorithm based on immersion simulations. IEEE Transactions on Pattern Analysis & Machine Intelligence 13, 583–598 (1991).

44. Yu, B. Stability. Bernoulli 19, 1484–1500 (2013).

45. Love, M. I., Huber, W. & Anders, S. Moderated estimation of fold change and dispersion for RNA-seq data with DESeq2. Genome Biol 15, 550 (2014).

